# Consciousness & Brain Functional Complexity in Propofol Anaesthesia

**DOI:** 10.1101/680447

**Authors:** TF Varley, A Luppi, I Pappas, L Naci, R Adapa, A Owen, DK Menon, EA Stamatakis

## Abstract

The brain is possibly the most complex system known to mankind, and its complexity has been called upon to explain the emergence of consciousness. However, complexity can take many forms: here, we investigate measures of algorithmic and process complexity in both the temporal and topological dimension, testing them on functional MRI data obtained from individuals undergoing various levels of sedation with the anaesthetic agent propofol, in two separate datasets. We demonstrate that the various measures are differently able to discriminate between levels of sedation, with temporal measures showing higher sensitivity. Further, we show that all measures are strongly related to a single underlying construct explaining most of the variance, as assessed by Principal Component Analysis, which we interpret as a measure of overall complexity of our data. This overall complexity was also able to discriminate between levels of sedation, supporting the hypothesis that consciousness is related to complexity - independent of how the latter is measured.

## 1 Introduction

The science of complex systems has gained increasing prominence in the 21st century. It combines the reductionist ideal of science, with the notion of emergence, whereby high-level phenomena can result from the interactions of simple constituent parts, confirming Aristotle’s saying that the whole is more than the sum of its parts [1]. However, complexity science is also a discipline still in its infancy. In particular, due to its appealing and apparently intuitive nature, the notion of complexity has remained relatively ill-defined. The interdisciplinary nature of this science has resulted in different fields applying the term complexity to multiple quantities, variously measured. Complexity is perhaps best understood as the negation of simplicity. A system exhibits complex behaviour when it is not uniform, stereotyped, or predictable. However, there is a key assumption that this is not sufficient: complexity must emerge from the underlying orderly interactions of a system’s components, about which its behaviour must provide information in other words, its unpredictability must be more than mere randomness, but rather the result of interesting behaviours emerging. Thus, a complex system lies between complete order such as the perfectly predictable regularity of a crystal and complete disorder, as exhibited for instance by the random motion of molecules of a gas. Complexity can be identified in more than one dimension of the same system, too. It may be due to the structure of the interactions between components, such as the connections in a social or biological network. Or it may only become apparent over time, as when it is applied to signals and temporal patterns. Furthermore, there are different ways in which something can be said to be complex, reflected in the different ways that have been developed to estimate complexity. On the one hand, methods from algorithmic information theory such as Shannon entropy and Lempel-Ziv compressibility [2, 3] emphasise unpredictability as the key property for complexity. One downside of such approach, however, is that they would treat a purely random sequence as maximally complex. Alternatively, methods from the physics of dynamical systems focus on the aspect of interactions in the process whether between the system’s elements (e.g. synchronisability; [4]), between its present and past states (e.g. Hurst exponent; [5]), or between different scales [6]. In this work, we aim to explore the relation between algorithmic and process measures of complexity, in both the topological and temporal dimensions. We choose to test these measures on a paradigmatically complex system: the human brain. Not only is the brain the source of humans’ widely diverse range of behaviours and accomplishments, which is itself suggestive of a highly complex underlying organisation; its structure is also that of a complex network of subnetworks, in turn made of multiple kinds of neurons obeying nontrivial plasticity rules for their interactions. For these reasons, it has been proposed that the brain’s complexity may explain another unique property it possesses: consciousness. Recent scientific theories of consciousness have emphasised, in one way or another, the brain’s complexity as a crucial requirement for consciousness [7, 8, 9, 10]. Anaesthetic drugs such as the GABA-ergic agonist propofol provide a way to controllably and reversibly modulate the brain’s state of consciousness. Its complexity, in various aspects, may then be assessed based on signals from noninvasive neuroimaging techniques. In particular, functional MRI (fMRI) has the advantage of providing high spatial resolution, thus allowing for estimation of the brain’s network properties in greater detail than afforded by other methods such as EEG. Here, we chose to evaluate measures of algorithmic and process complexity applied to the temporal and topological (network) dimensions, derived from fMRI blood-oxygen-level-dependent (BOLD) signals of volunteers undergoing sedation with propofol, in order to investigate the relationship between the different measures of complexity, as well as determining whether they can be related to different levels of consciousness. We also replicated our results in an independent dataset of propofol anaesthesia, in order to demonstrate their robustness.

## 2 Methods

### 2.1 Data Acquisition & Preprocessing

#### 2.1.1 Dataset A

Twenty-five healthy volunteer subjects were recruited for scanning. The acquisition procedures are described in detail by Stamatakis et al, [11]: MRI data were acquired on a Siemens Trio 3T scanner (WBIC, Cambridge). Each functional BOLD volume consisted of 32 interleaved, descending, oblique axial slices, 3 mm thick with interslice gap of 0.75 mm and in-plane resolution of 3 mm, field of view = 1926192 mm, repetition time = 2 s, acquisition time = 2 s, time echo = 30 ms, and flip angle 78. We also acquired T1-weighted structural images at 1 mm isotropic resolution in the sagittal plane, using an MPRAGE sequence with TR = 2250 ms, TI = 900 ms, TE = 2.99 ms and flip angle = 9u, for localization purposes. Of the 25 healthy subjects, 14 were ultimately retained: the rest were excluded, either because of missing scans (n=2), or due of excessive motion in the scanner (n=9, 5mm maximum motion threshold).

##### Propofol Sedation

Propofol was administered intravenously as a target controlled infusion (plasma concentration mode), using an Alaris PK infusion pump (Carefusion, Basingstoke, UK). Three target plasma levels were used - no drug (baseline), 0.6 mg/ml (mild sedation) and 1.2 mg/ml (moderate sedation). A period of 10 min was allowed for equilibration of plasma and effect-site propofol concentrations. Blood samples were drawn towards the end of each titration period and before the plasma target was altered, to assess plasma propofol levels. In total, 6 blood samples were drawn during the study. The mean (SD) measured plasma propofol concentration was 304.8 (141.1) ng/ml during light sedation, 723.3 (320.5) ng/ml during moderate sedation and 275.8 (75.42) ng/ml during recovery. Mean (SD) total mass of propofol administered was 210.15 (33.17) mg, equivalent to 3.0 (0.47) mg/kg. The level of sedation was assessed verbally immediately before and after each of the scanning runs. The three conditions from this dataset are referred to as Awake, Mild and Moderate sedation respectively.

#### 2.1.2 Dataset B

These data were generously provided by the Brain and Mind Institute, Department of Psychology, The University of Western Ontario. Sixteen healthy volunteer subjects were recruited for scanning. Scanning was performed using a 3 Tesla Siemens Tim Trio system with a 32-channel head coil, at the Robarts Research Institute in London, Ontario, Canada. Participants lay supine in the scanner. Function echo-planar images (EPI) were acquired (33 slices, voxel size: 3 x 3 x 3mm; inter-slice gap of 25%, TR=2000ms, TE=30ms, matrix size=64×64, FA=75 degrees). An anatomical volume was obtained using a T1-weighted 3D MPRAGE sequence (32 channel coil, voxel size: 1x 1 x 1mm, TA= 5 min, TE=4.25ms, matrix size=240×256, FA=9 degrees).

##### Propofol Sedation

Intravenous propofol was administered with a Baxter AS 50 (Singapore). The infusion pump was manually adjusted using step-wise increases to achieve desired levels of sedation of propofol (Ramsay level 5). Concentrations of intra-venous propofol were estimated using the TIVA Trainer (the European Society for Intravenous Aneaesthesia, eurosiva.eu) pharmacokinetic simulation program. If Ramsay level was lower than 5, the concentration was slowly increased by increments of 0.3 *µ*g/ml with repeated assessments of responsiveness between increments to obtain a Ramsay score of 5. Ramsay level 5 was determined as being unresponsive to verbal commands and rousable only by physical stimulus. In contrast to Propofol Dataset A, the two conditions from this dataset are referred to by Awake and Deep sedation respectively, reflecting the substantial increase in sedation depth present in this dataset.

#### 2.1.3 Image Pre-Processing

All of the collected images were preprocessed using the CONN functional connectivity toolbox [12]^1^, using the default pre-processing pipeline, which includes realignment and unwarping (motion estimation and correction), slice-timing correction, outlier detection, structural coregistration and spatial normalisation using standard grey and white matter masks, normalization to the Montreal Neurological Institute space (MNI), and finally spatial smoothing with a 6mm full width at half-maximum (FWHM) Gaussian kernel.

Temporal preprocessing included nuisance regression using anatomical CompCor to remove noise attributable to white matter and CSF components from the BOLD signal, as well as subject-specific realignment parameters (three rotations and three translations) and their first-order temporal derivatives [13]. Linear detrending was also applied, as well as band-pass filtering in the default range of [0.008, 0.09] Hz [14]. For a more detailed discussion of the details of the CONN default preprocessing pipeline, see Whitefield-Gabrieli and Nieto-Castanon, 2012.

### 2.2 Complexity of BOLD Signals

To explore the space of different formalizations of complexity, we used algorithms from algorithmic information theory (Lempel-Ziv compressibility, sample entropy, and principal component analysis), as well as from dynamical systems physics (Higuchi fractal dimension, Hurst exponent). Before analysis, the BOLD time-series were transformed by applying the Hilbert transform. The absolute value of the transformed signal was then taken, to remove negative frequencies and ensure that all series were positive. The Hilbert transform was also used to maintain consistency with earlier studies exploring the complexity of brain activity as it relates to consciousness [15, 16].

#### 2.2.1 Lempel-Ziv Complexity

The Lempel-Ziv algorithm is a computationally tractable method for quantifying the complexity of a data-series by calculating the number of distinct patterns present in the data. For sufficiently large datasets, it is a useful approximation of Kolmogorov complexity, which is famously uncomputable for most strings [2]. The method used here is described in Shartner et al., (2015). Briefly: for every ROI in our parcellated brain, a time-series *F*(*t*) is binarized according to the following procedure:

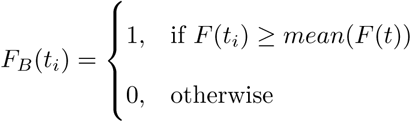

The resulting time-series are stacked into a binary matrix *M*(*X, T*), where every row corresponds to the time-series *F*_*B*_(*t*) for every ROI *x* ∈ *X* and every column is a time-point *t* ∈ *T*. The matrix is then flattened orthogonally to *T*, resulting in a vector *V* of length *X* × *T*, on which the Lempel-Ziv analysis was performed.

The Lempel-Ziv algorithm creates a dictionary *D*, which is the set of binary patterns that make up *V* and returns a value *LZ*_*C*_ ∝ |*D*|. For every time-series *F*_*B*_(*t*) ∈ *X*, a random time-series was created, by shuffling all the entries in *F* (*t*). These were stacked into a binary matrix *M*_*rand*_, with the same dimensions as *M*, however containing only noise. This random matrix was flattened and its *LZ*_*C*_ value calculated. As the randomness of a string increases, *LZ*_*C*_ → 1, so this value was used to normalize the “true” value of *LC*_*C*_, which was divided by 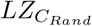 to ensure all values were within a range (0, 1).

#### 2.2.2 Sample Entropy

Sample Entropy (SampEn) quantifies how unpredictable a signal is [3] by estimating the probability that similar sequences of observations in a timeseries will remain similar over time. To compute SampEn, each time-series *X*(*t*) of length *N* is divided into subsections *S* of length *m* and the Chebychev distance between two sections *S*_*i*_, *S*_*j*_ is calculated. Two sections are “similar” if their distance is less than some tolerance *r*. The procedure is repeated for sections of length *m* + 1. We then calculate the probability that, if two data sequences of length *m* have distance less than *r*, then the same two sequences of length *m* + 1 also have distance less than *r*.

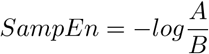

Where A is the number of chunks of length *m* + 1 that are similar (have Chebyshev distance less than *r*), and B is the number of chunks of length *m* that are similar. Low values of SampEn would indicate that the signal is highly stereotyped - with a perfectly predictable series, such as [1,1,1, …] having a SampEn of zero, and SampEn increasing as the series becomes more disordered.

SampEn depends on the choice of parameters *m* and *r*. Here, we used *m* = 2 and *r* = 0.3 × *σ*(*X*(*t*)), where *σ*() is the standard deviation function.

SampEn has been used to test the level of sedation induced by propofol and remifentanil in electrophysiological studies (Ferenets et al., 2007), and been shown to be associated with the degree of sedation much like Lempel-Ziv complexity has.

#### 2.2.3 Hurst Exponent

The Hurst Exponent returns an estimate of how predictable a time-series is by quantifying its ‘memory,’ or how dependent the value at time *t* is on the value at time *t* − 1 [5]. There are a number of algorithms for estimating the Hurst Exponent; here we report results calculated using a rescaled range approach. In it, a time-series *X*(*t*) of length *N* is segmented into non-overlapping sections of length *n, X*_*i*_(*t*). For each segment, the cumulative departure from the signal mean is calculated:

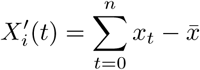

where 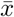 is the mean of *X*_*i*_(*t*). The rescaled range of deviations (*R/S*) is then defined as:

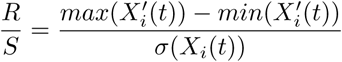

where *σ*() is the standard deviation function. We then compute *R/S* for all *X*_*i*_(*t*) and average them, generating (*R*(*n*)*/S*(*n*)), which is the average scaled range for all the subsections of *X*(*t*) with length *n*. We are left with a power relation, where:

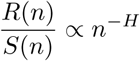

Where *H* is the Hurst exponent, and can be extracted by regression.

#### 2.2.4 Higuchi Fractal Dimension

To calculate the temporal fractal dimension, we used the Higuchi method for calculating the self-similarity of a one-dimensional time-series [6], an algorithm widely used in EEG and MEG analysis [17]. From each time-series *X*(*t*), we create a new time-series 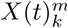, defined as follows:

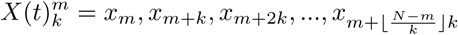

where *m* = 1, 2, …, *k*.

For each time-series 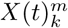 in *k*_1_,*k*_2_, …*k*_*max*_, the length of that series, *L*_*m*_(*k*), is given by:

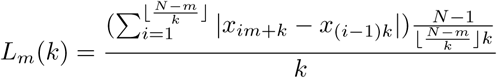

We then define the average length of the series ⟨*L*(*k*)⟩, on the interval [*k, L*_*m*_(*k*)] as:

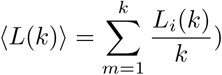

If our initial time-series *X*(*t*) has fractal character, then:

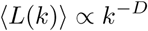

Where *D* is our desired fractal dimension. The Higuchi algorithm requires a pre-defined *k*_*max*_ value as an input, along with the target time-series. This value is usually determined by sampling the results returned by different values of *k*_*max*_ and selecting a value based on the range of *k*_*max*_ where the fractal dimension is stable. For both datasets, we sampled over a range of powers of two (2, …, 128). Due to the comparably small size of BOLD time-series, the range of *k*_*max*_ values that our algorithm could process without returning an error was limited. We ultimately decided on *k*_*max*_ = 32 for Dataset A and *k*_*max*_ = 64 for the Dataset B.

#### 2.2.5 PCA of BOLD Signals

Principal component analysis (PCA) is commonly used to compress data by finding the dimensions that encode the maximal variance in a high-dimensional dataset. Here, we use PCA in a matter similar to Lempel-Ziv complexity, to relate the complexity of sets of BOLD signals to their compressibility. The more algorithmically random the dataset, the more orthogonal dimensions are required to describe the dataset, which we took advantage of to attempt to quantify the complexity of our BOLD time-series data. We constructed a large array of un-binarized BOLD signals, *M*(*X, T*) to which we applied a standard feature scaler from Scikit-Learn [18] to ensure all values had a mean of zero and unit variance, and then a PCA function, recording recorded how many dimensions were required to cumulatively describe 95% of the variance in the original dataset. We used this value as our measure of data complexity.

### 2.3 Complexity of Functional Connectivity Graphs

Networks are a common example of complex system, and perhaps none more so than the human brain, which can be considered as a network at multiple scales. A network, or graph, is represented mathematically as an object comprised of nodes (in this case, cortical regions) and the connections between them, or edges (in this case, functional connectivity given by statistical association of the regions’ BOLD time-series). Investigating how the complexity of brain functional networks is affected by the anaesthetic drug propofol is therefore a clear way of testing our hypothesis that loss of consciousness should reduce the brain’s level of complexity.

#### 2.3.1 Formation of Functional Connectivity Networks

To construct brain functional connectivity networks, the preprocessed BOLD time-series data were extracted from each brain in CONN and the cerebral cortex was segmented into distinct ROIs, using the 234-ROI parcellation of the Lausanne atlas [19]. Each time-series *F*(*t*) was transformed by taking the norm of the Hilbert transform, to maintain consistency with the time-series analysis.

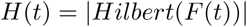

Every time-series *H*(*t*) was then correlated against every other time-series, using the Pearson Correlation, forming a matrix *M* such that:

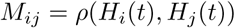

The matrices were then filtered to remove self-loops, ensuring simple graphs, and all negative correlations were removed:

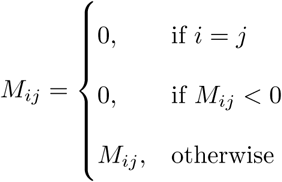

Finally, the matrices were binarized with a *k*% threshold, such that:

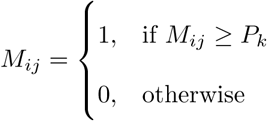

The results could then be treated as adjacency matrices defining functional connectivity graphs, where each row *M*_*i*_ and column *M*_*j*_ corresponds to an ROI in the initial cortical parcellation, and their connection being represented by the corresponding cell in the matrix. For each graph theoretical analysis, a range of percentage thresholds (*k*%) were tested to ensure that any observed effects were not an artefact of one particular threshold, and are consistent over different graph topologies.

#### 2.3.2 Algebraic Connectivity

Algebraic connectivity (AC) is a measure of graph connectivity derived from spectral graph theory [4], which gives an upper bound on the classical connectivity of a graph. As such, it is often used as a measure of how well-integrated a graph is and how robust it is to damage, in the sense of the number of connections that must be removed before it is rendered disconnected. Unlike classical connectivity, which must be calculated by computationally intensive brute-force methods, AC is quite easy to find for even quite large graphs. AC is also a measure of graph synchronizability and emerges from analysis of the Kuramoto model of coupled oscillators [20]. For a simple example, imagine placing identical metronomes at every vertex of a graph and allowing the vibrations to propagate along the edges. The synchronizability describes the limit behaviour of how long it will take all the metronomes to synchronize. Here we use AC as a proxy measure of synchronisability to capture the possible temporal dynamics of the brain networks modelled by our functional connectivity graphs.

The AC of a graph *G* is formally defined as the first non-zero eigenvalue of the Laplacian matrix *L*_*G*_ associated with *G. L*_*G*_ is derived by subtracting the adjacency matrix *A*_*G*_ from the degree matrix *D*_*G*_:

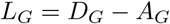

As every row and column of *L*_*G*_ sum to zero, and it is symmetric about the diagonal, the imaginary part of every eigenvalue in the spectrum of *L*_*G*_ is zero, and if *G* is a fully-connected graph, then:

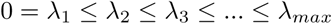

To ensure that we were capturing the full topology of the graph, we calculated λ_2_ for each graph at multiple thresholds [10, 20, 30, … 90], creating a curve 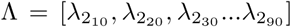. We then integrated Λ using the trapezoid method to arrive at our final value *AC* = ∫ Λ *dx*.

#### 2.3.3 Graph Compressibility

In contrast to AC, which we use to explore the limit behaviour of possible brain temporal dynamics, our measure of graph compressibility is purely algorithmic, and estimates the Kolmogorov complexity of a graph: that is, the size of a computer program necessary to fully recreate a given graph *G*. To do this, we re-employ the Lempel-Ziv algorithm originally used to calculate the *LZ*_*C*_ score of BOLD signals. Here we use it to calculate a related measure, *LZ*_*G*_, which is the length of a dictionary required to describe the adjacency matrix *A*_*G*_ of a given graph.

To calculate *LZ*_*G*_, we take a binary adjacency matrix and flatten it into a single vector *V*, and then run the Lempel-Ziv algorithm on that vector. As a binary vector of length *l* can be used to perfectly reconstruct an adjacency matrix defining a graph with 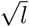 vertices (so long as *l* is a square number, of course), we take *V* to be equivalent to a program defining *G*. As with AC, to ensure that we were capturing the full topology of *G*, we calculated the Lempel-Ziv complexity of the binary *A*_*G*_ at the same nine thresholds [10…90], and then defined *LZ*_*G*_ as the integral of the resulting curve of complexity values.

### 2.4 Higher-Order Measures

Once we had calculated individual measures of complexity, we tested how they related to each-other, and (for Dataset A) serum concentrations of propofol. We correlated each one against all others to construct a correlation matrix which describes, how different metrics cluster.

We also did a principal component analysis on the set of all results. We hypothesized that, despite variability in the effectiveness of the individual measures, there should be a single, underlying component reflecting a shared factor of “complexity”. We further hypothesised that this underlying factor should be predictive of both the level of consciousness, and (in Dataset A), of the individual serum concentration of propofol.

### 2.5 Statistical Analysis

All analysis was carried out using the Python 3.6 programming language in the Spyder IDE ^2^, using the packages provided by the Anaconda distribution ^3^. All packages were in the most up-to-date version. Packages used include SciKit-Learn [18], NumPy [21], SciPy [22], and NetworkX [23]. Unless otherwise specified, all the significance tests are non-parametric: given the small sample sizes and heterogeneous populations, normal distributions were not assumed. Wilcoxon Signed Rank test was used to compare drug conditions against their respective control conditions.

## 3 Results

### 3.1 Temporal Algorithmic Complexity

#### 3.1.1 Lempel-Ziv Compressibility

The first measure of algorithmic complexity we used was normalized Lempel-Ziv compressibility [15, 16] of BOLD signals. We found significant differences between conditions in both Dataset A and Dataset B. In Dataset A, Kruskal-Wallis Analysis of Variance found significant differences between all three conditions (H(10.57), p=0.005), and post-hoc analysis with Wilcoxon Signed-Rank test found significant differences between the Awake and Mild conditions (W(21), p=0.05), Awake and Moderate conditions (W(9), p=0.006), and Mild and Moderate conditions (W(4), p=0.002). In Dataset B, the Wilcoxon test found significant difference between the Awake and Deep conditions (H(19), p=0.011). In both datasets, the Awake condition had the highest complexity, and as the depth of sedation increased, the associated LZC decreased. In Dataset A, the transition from Awake to Moderate showed Δ = −0.029 ± 0.027, and in Dataset B, the transition from Awake to Deep showed Δ =−0.029 ± 0.039. We note that these two results are remarkably similar, although this is likely a coincidencee. For full results from Dataset A, see Table 1, and for Dataset B, Table 2. In the propofol sedation conditions of Dataset A (Mild and Moderate), we found significant negative correlations between LZC and serum concentrations of propofol (r=-0.55, p=0.002), see Figure 3A.

**Table 1:** The values for all the complexity measures, temporal and spatial, for Dataset A.

| | $LZ_C$ | SampEnt | PCA | Hurst | Higuchi | AlgConn | $LZ_{Graph}$ | Serum Propofol |
| --- | --- | --- | --- | --- | --- | --- | --- | --- |
| Awake | $0.967 \pm 0.013$ | $0.662 \pm 0.013$ | $28.5 \pm 7.771$ | $0.764 \pm 0.009$ | $0.867 \pm 0.015$ | $470.433 \pm 73.893$ | $375316.786 \pm 4256.981$ | N/A |
| Mild | $0.962 \pm 0.012$ | $0.654 \pm 0.01$ | $27.429 \pm 7.5$ | $0.769 \pm 0.008$ | $0.864 \pm 0.013$ | $436.068 \pm 70.707$ | $375353.214 \pm 3931.755$ | $286.025 \pm 133.5$ |
| Moderate | $0.939 \pm 0.028$ | $0.626 \pm 0.035$ | $26.286 \pm 7.314$ | $0.778 \pm 0.013$ | $0.842 \pm 0.025$ | $362.762 \pm 85.815$ | $372431.071 \pm 4277.4$ | $626.126 \pm 249.8$ |

**Table 2:** The values for all the complexity measures, temporal and spatial, for Dataset B.

| | $LZ_C$ | SampEnt | PCA | Hurst | Higuchi | AlgConn | $LZ_{Graph}$ |
| --- | --- | --- | --- | --- | --- | --- | --- |
| Awake | $0.967 \pm 0.018$ | $0.659 \pm 0.017$ | $45.812 \pm 2.744$ | $0.738 \pm 0.008$ | $0.981 \pm 0.011$ | $4693.704 \pm 826.809$ | $372236.562 \pm 3794.728$ |
| Deep | $0.938 \pm 0.05$ | $0.636 \pm 0.038$ | $40.625 \pm 4.702$ | $0.749 \pm 0.016$ | $0.963 \pm 0.03$ | $3787.616 \pm 1338.374$ | $367313.125 \pm 8565.381$ |

**Figure 1:**
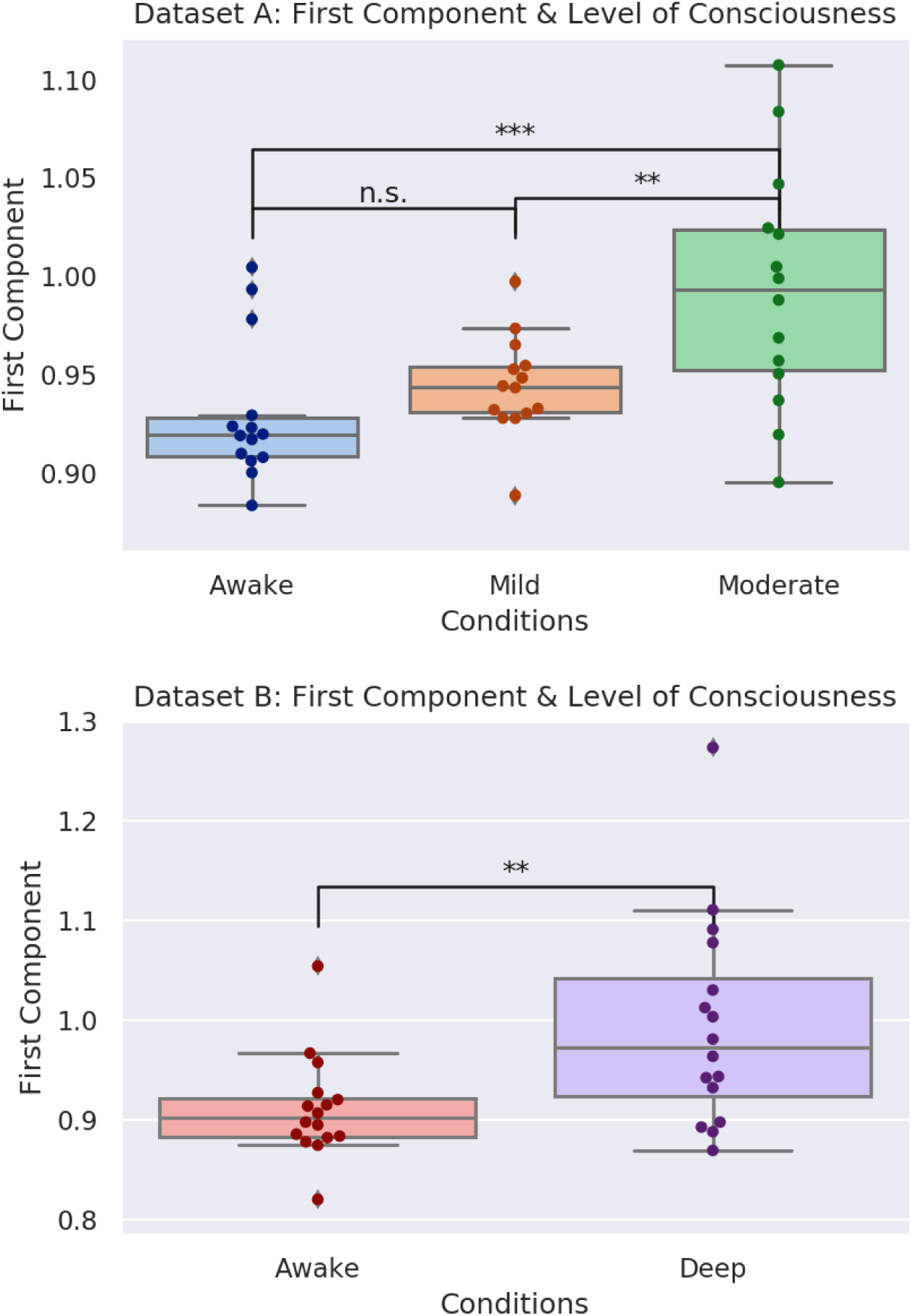
Here are the differences in the first principal component generated from all the measures from Datasets A and B. Interestingly, in Dataset A, there was no significant difference between the Awake and Mild condition, while there were differences between both of those states and the Moderate condition. While this may be a reflection of lack of sensitivity, it is worth noting that, between the Awake and Mild conditions, consciousness was not actually lost: volunteers experienced conscious sedation, while the difference in level of consciousness between the Awake and and Moderate conditions was much more dramatic. In Dataset B, where consciousness was fully lost in the Deep condition, a significant difference appeared. Note that, despite the measures of complexity generally dropping as consciousness was lost (with the notable exception of the Hurst exponent analysis), the PCA analysis returned a Hurst-like pattern, with the values in the component increasing as consciousness is lost. This does not indicate an increase in complexity in any sense, but rather, is an artefact of how the dimensionality reduction transforms values. To ensure that this was not being driven by the Hurst exponent in any way, we ran the analysis after multiplying each Hurst exponent by −1 (so that the value decreased with loss of co2n9sciousness), and found no difference in the result.

**Figure 2:**
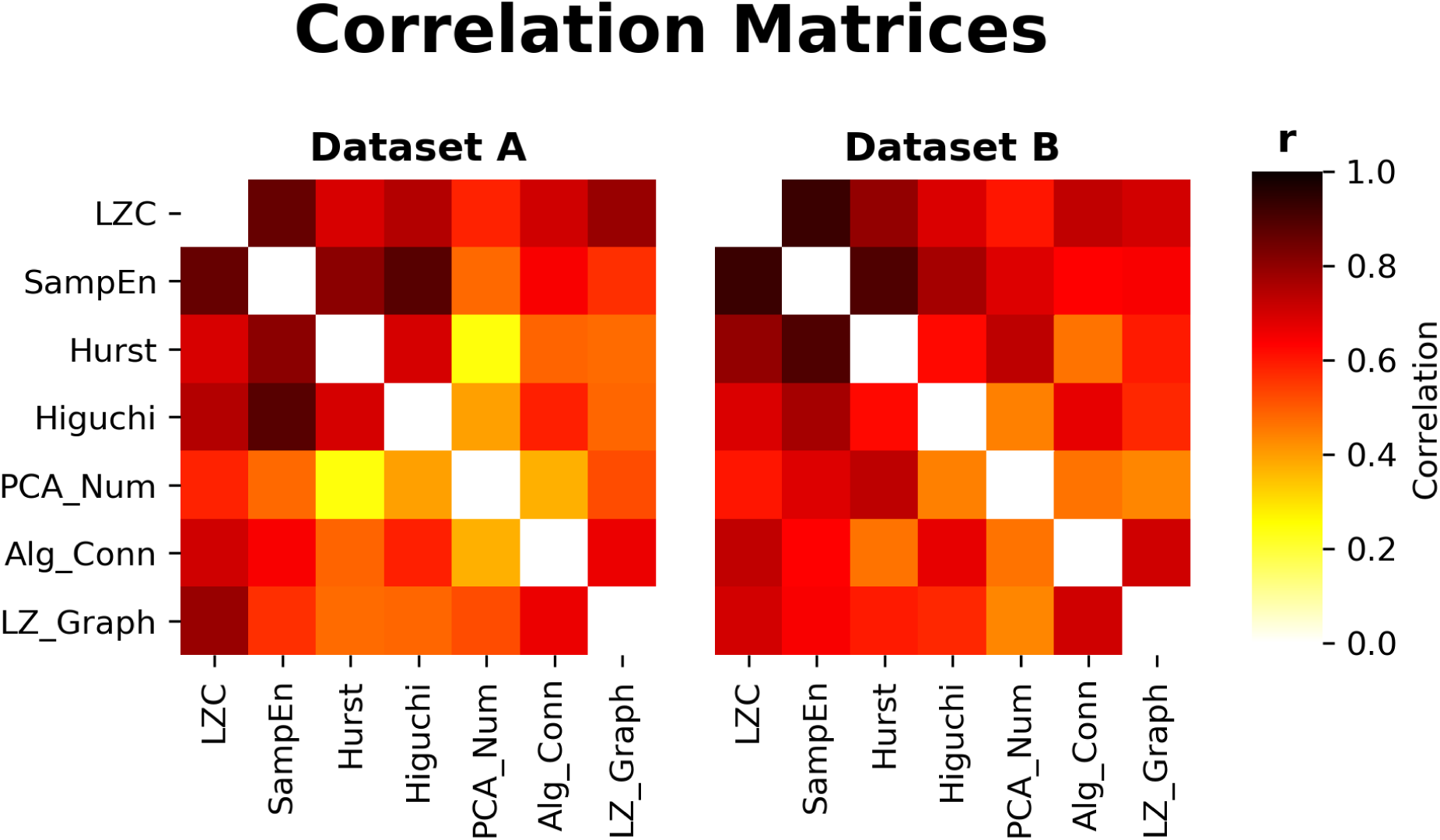
The correlation matrices between all the different metrics fro Datasets A and B. All entries along the diagonal have been removed. There are some typical patterns: the graph measures (LZ Graph and Algebraic Connectivity are both generally more highly correalted, as are LZC, Sampen and Hurst). With the exception of a single correlation between the PCA Number and the Hurst Exponent in Dataset A. The p-values ranged over many orders of magnitude from 10^−2^ to 10^−20^

**Figure 3:**
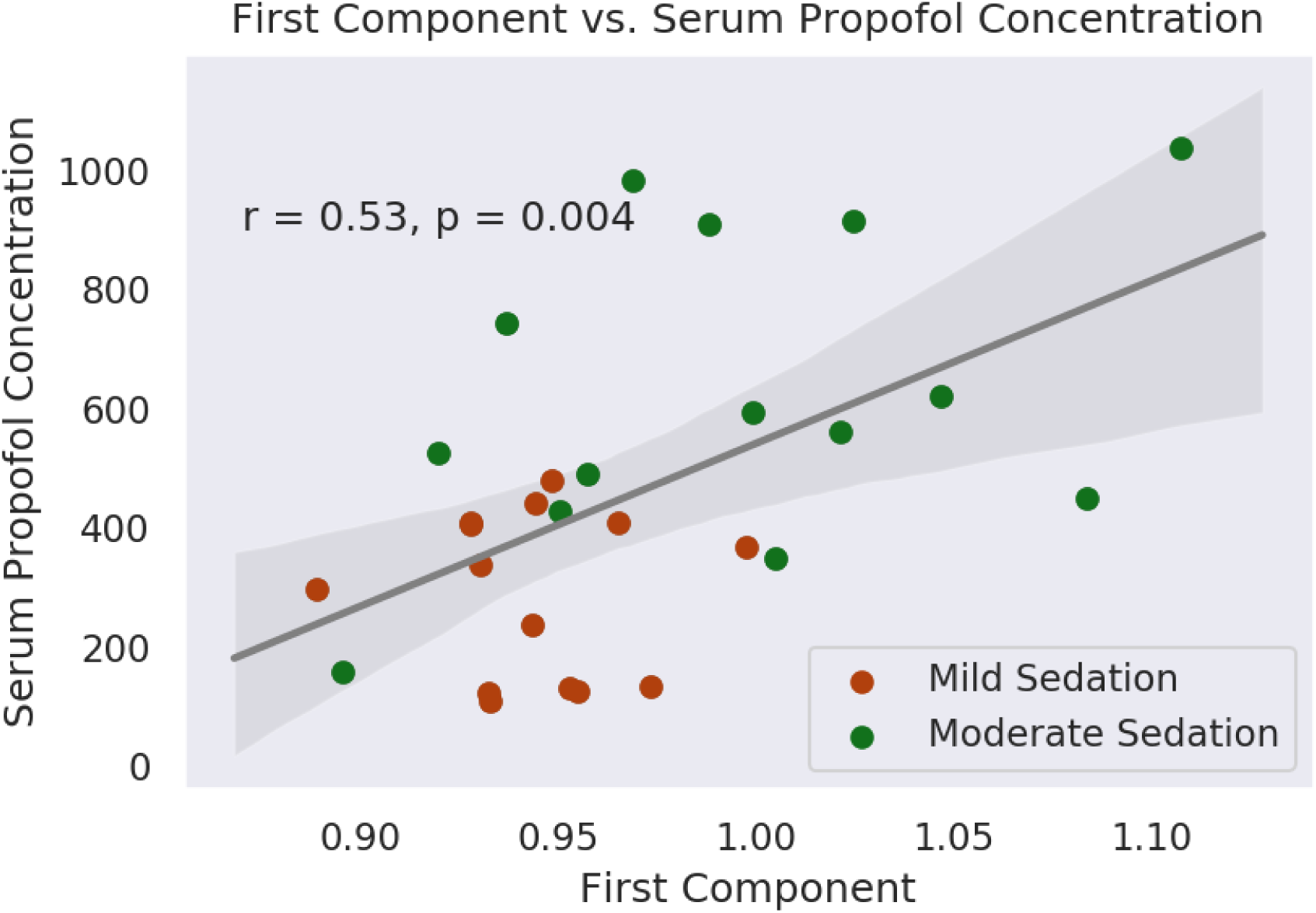
There was a significant correlation between the first component and serum concentration of propofol, with patients in the Mild condition (r = 0.53, p-value = 0.004) clustering together with low concentrations, and increasing, with larger variances, as the propofol concentration climbs. As with the plots shown above, the incongruous increase in the values of the component does not reflect a relative increase in complexity in this case, but is an artefact of the PCA algorithm. No Awake volunteers were included in this analysis, as all would have had a blood propofol concentration of exactly zero.

These results are consistent with previous findings that Lempel-Ziv compressibility of spontaneous brain activity is discriminative of level of consciousness in humans [15, 16] and animals [24].

Of all the time-series measures described, the LZC algorithm described here is distinct in that it communicates information about the spatial complexity as well as the temporal complexity. This is because, unlike other measures like Sample Entropy or Higuchi Fractal Dimension which are calculated on 234 individual time-series and then averaged, LZC is calculated on an entire dataset, which has been flattened column-wise, as was done in [15, 16, 25], by “stacking” each column on top of the next, resulting in a one-dimensional vector where the first 234 elements are the first column, the second 234 elements are the second column, etc. This means that the vector *V* (see Methods section) can be divided into 234 segments where every entry corresponds to the coarse activation of a distinct brain region at the same time. The result is that each entry in the dictionary *D* created by the Lempel-Ziv algorithm corresponds, not to a series of samples from a single ROI, but rather a distribution of cortical regions that are “on” or “off.”

#### 3.1.2 Sample Entropy

We found significant decreases in the Sample Entropy of BOLD signals under anaesthesia in both Datasets A and B. In Dataset A, Kruskal-Wallis Analysis of Variance found a significant difference between all three conditions (H(12.94), p=0.002) and post-hoc analysis with the Wilcoxon Signed-Rank test found significant differences between the Awake and Moderate conditions (W(6), p=0.004), and Mild versus Moderate conditions (W(8), p=0.005), but not the Awake versus Mild conditions. In Dataset B there was a significant difference between the Awake and Deep conditions (W(21), p=0.015). As with the LZC analysis, the Awake condition had the highest Sample Entropy in both Datasets A and B, with the mean value decreasing with increasing sedation. In Dataset A, we observed Δ = 0.036 ± 0.035 from Awake to Moderate, and in Dataset B we observed Δ = −0.023 ± 0.031. In the Mild and Moderate conditions of Dataset A, we found a significant negative correlation between serum concentration of propofol and Sample Entropy of BOLD signals (r=-0.53, p=0.003).

These results are consistent with both the LZC results reported above and the findings of Ferentes et al., (2007), who found that Sample Entropy decreased with increasing sedation in much the same way that LZC does.

#### 3.1.3 PCA of BOLD Signals

As with LZC, the PCA-based measure of BOLD signal complexity returns a measure of how compressible the set of data are as a proxy for complexity, by identifying the number of components required to explain a fixed proportion of the variance in the data. A larger number of components to explain the same amount of variance would indicate less compressibility of the data. Thus, we hypothesized that as level of sedation increased, so would the compressibility of BOLD signals, as measured by the number of components required to explain 95% of the variance. In Dataset A, Kruskal-Wallis analysis of variance found significant differences between all three conditions (H(8.13), p=0.017), and post-hoc testing found significant differences between all three sets of conditions: Awake vs. Mild (W(9), p=0.03), Awake vs. Moderate (W(6), p=0.016), and Mild vs. Moderate (W(11), p=0.048). In Dataset B, we found a significant difference between Awake and Deep (W(4), p=0.002). As before, there was a consistent pattern of increasing mean compressibility (and a consequent decreasing number of required components) as sedation increased. In Dataset A, Δ = −2.214 ± 2.73 from Awake to Moderate, and in Dataset B, Δ = −5.188 ± 4.68. Here, the Δ is negative because the number of components decreased between the Awake and sedated conditions, and is non-integer because it is the average over all subjects in the datasets. Of all the measures of BOLD signal compressibility, this was the only measure that did not significantly correlate with serum propofol concentration in the Mild and Moderate conditions in Dataset A.

As with LZC and the SampEn, these results indicate that as propofol-induced sedation increases, the algorithmic complexity of BOLD signals decreases. All measures of complexity discussed so far support each-other, despite being a variety of linear and non-linear algorithms.

### 3.2 Temporal Process Complexity

#### 3.2.1 Hurst Exponent

The Hurst Exponent was the only measure that we hypothesized would increase as consciousness was lost, rather than decrease, since as a signal becomes more predictable, its Hurst Exponent tends towards unity [5]. In both Datasets A and B we found significant differences between conditions. In Dataset A, Kruskal-Wallis Analysis of Variance found an omnibus difference (H(9.11), p=0.01), and post-hoc testing found significant differences between Awake and Mild (W(16), p=0.022), Awake and Moderate (W(8), p=0.005), and Mild and Moderate (W(17), p=0.026). In Dataset B, we found significant differences between the Awake and Deep conditions (W(26), p=0.02). Unlike the previous two metrics, and as we expected, we found a relative increase in the Hurst Exponent as sedation increased: in Dataset A, we found Δ = 0.014 ± 0.014 from Awake to Moderate sedation, and in Dataset B we found Δ = 0.01 ± 0.016 from Awake to Deep sedation. In Dataset A, we found a significant correlation between serum concentration of propofol and Hurst Exponent in the Mild and Moderate sedation conditions (r=0.393, p=0.039).

This is consistent with our initial hypothesis that as sedation increased and consciousness was lost, the BOLD signals would become more predictable, as measured by an increasing Hurst Exponent.

#### 3.2.2 Higuchi Fractal Dimension

The Higuchi Fractal Dimension was one of the least sensitive measures of BOLD signal complexity sampled here. In Dataset A, Kruskal-Wallis Analysis of Variance found a significant difference between all three conditions (H(8.27), p=0.016), and post-hoc analysis found significant differences between the Awake and Moderate conditions (W(15), p=0.019) and the Mild and Moderate conditions (W(17), p=0.026), but not the Awake and Mild conditions. In Dataset B we found a significant difference between the Awake and Deep conditions (W(23), p=0.02). As with LZC and Sample Entropy, the Awake condition had the highest mean fractal dimension in both samples, which went down as sedation increased: in Dataset A Δ = −0.024 ± 0.032 from Awake to Moderate and in Dataset B, Δ = −0.018 ± 0.027 from Awake to Deep. Surprisingly, the Higuchi Fractal dimension showed a very strong negative correlation with serum propofol concentration in the Mild and Moderate conditions of Dataset A (r=-0.614, p=0.0005).

The finding that Higuchi Fractal dimension was relatively less able to discriminate between level of consciousness than LZC or Sample Entropy but more predictive of serum propofol concentration is interesting. While it is hard to come up with a definitive interpretation, it may suggest that there is some variable factor in individuals that makes their level of consciousness more or less resistant to the changes in brain activity (as measured by Higuchi Fractal Dimension) induced by propofol, or that the plasma concentration data offer more resolution, extending beyond the three artificially imposed bins of Awake, Mild and Moderate sedation.

### 3.3 Topological Algorithmic Complexity

#### 3.3.1 Algebraic Connectivity

Our first of two measures of functional network complexity is algebraic connectivity, which returns information about the robustness of the network to removal of elements [26]. In Dataset A, Kruskal-Wallis analysis found a significant difference in algebraic connectivity between all three conditions (H(9.654), p=0.008). Post-hoc analysis found significant differences between the Awake and Moderate conditions (W(12), p=0.011) and the Mild and Moderate conditions (W(15), p=0.019), but not the Awake versus Mild conditions. In Dataset B, we found a significant difference between the Awake and Deep conditions (W(23), p=0.02). As before, in Datasets A and B the Awake condition had the highest mean algebraic connectivity, with mean values dropping as sedation increased. In Dataset A, Δ = −107.67 ± 116.98 from Awake to Moderate, while in Dataset B, −906.09 ± 1235.07. Despite the ability of algebraic connectivity to discriminate between conditions, there was no significant correlation with serum propofol concentration in the Mild and Moderate conditions of Dataset A.

These results suggest that, while graph theoretical measures may be predictive of level of consciousness in propofol anaesthesia, algebraic connectivity in particular seems to lack the discriminative power of direct analysis on BOLD signals. Nevertheless, these results are promising as they show that the topological complexity of functional brain networks can communicate information relevant to the level of consciousness of an individual.

### 3.4 Topological Process Complexity

#### 3.4.1 Graph Lempel-Ziv Compressibility

The final metric we tested, and the second measure of topological complexity, was the compressibility of functional connectivity adjacency matrices using the Lempel-Ziv algorithm. This was the weakest of all the measures explored: the only significant difference was in Dataset A, between the Awake and Moderate conditions (W(11), p=0.009), although in Dataset B there was a similar trend that approached, but did not reach significance (W(32), 0.06). The general trend of Awake having the highest value which decreased under increasing sedation was conserved (although note the large standard deviations): in Dataset A Δ = −2885.71 ± 4937.34 and in Dataset B Δ = −4923.44 ± 8967.29. There was no significant correlation between graph compressibility and serum propofol concentrations in Dataset A.

While this is clearly the weakest result, in the context of the others, we still find its success at discriminating between the Awake and Moderate conditions of Dataset A intriguing, and suspect that in a larger set of data it may have more discriminative power. The relationship between consciousness and network compressibility may not be as direct as when performing analysis such as LZC on BOLD signals, but these results suggest this is an area worth exploring.

### 3.5 Higher Order Analysis of Overall Complexity

Every metric, when correlated against every other metric, showed a highly significant correlation (see Figure 2), all of which were significant with the sole exception of the correlation between the number of PCA components required to explain the majority of the variance and the Hurst exponent in Dataset A. We had hypothesized that, if the different kinds of complexity explored here (algorithmic and process-based, in both the temporal and topological dimensions) all were ways to quantify an underlying construct of overall complexity, then there should be a single component that explains the majority of the variance of the results. In Dataset A, we found that the principal component explained 67.07% of the variance in the set of results and in Dataset B the principal component explained 71.05% of the variance of the results. In both datasets, this component correlated extremely highly with each metric: in Dataset A it correlated most highly with LZC (r=-0.947, p≤ 1 × 10^−5^), followed by Sample Entropy (r=-0.929, p≤ 1 × 10^−5^). In Dataset B, these two were also the most highly correlated with the principal component, although the order was flipped, with Sample Entropy having the highest correlation (r=-0.95, p≤ 1 × 10^−5^), followed by LZC (r=-0.932, p≤ 1 × 10^−5^). When broken down by condition, in both Datasets, the principal component was able to discriminate between states of consciousness: in Dataset A the Kruskal-Wallis test found a significant difference between all three conditions (H(12.048, p=0.002), and post-hoc testing found significant differences between the Awake and Moderate conditions (W(8), p=0.005), and the Mild and Moderate conditions (W(3), p=0.001) but not the Awake and Mild conditions. In Dataset B we found a significant difference between the Awake and Deep conditions (W(8), p=0.002). In the Mild and Moderate conditions of Dataset A, the principal component significantly correlated with serum concentrations of propofol (r=0.531, p=0.004). Thus, the principal component derived from multiple specific measures of complexity can be related to states of consciousness in the human brain, and may be identified with the overall complexity of the dataset.

## 4 Discussion

In the present work, we have investigated measures of complexity from algorithmic information theory and the physics of dynamical systems, as they apply to the temporal and topological (network) dimensions of functional MRI brain data from individuals under different levels of propofol sedation. Two main insights can be derived from our results. The first is that, at least in the context of the human brain, different measures purporting to quantify complexity are indeed related to some underlying common construct, regardless of the dimension along which they measure complexity, or the aspect of complexity that they measure. This provides much-needed validation to the idea that a dataset - and the system from which it derives - can be considered complex *tout court*, rather than just being complex in a specific dimension, and according to a specific way of assessing complexity. We term this the overall complexity of the system or dataset. In turn, this suggests that it is appropriate to use the term complexity for the various specific measures, because there does seem to exist a common underlying property of the data that they tap into. In particular, we have demonstrated that the complexity of the human brain activity, as inferred from fMRI BOLD signals, is modulated by one’s state of consciousness. This was observed both with the individual measures - validating and extending previous results - and, most importantly, with the underlying construct of overall complexity, which demonstrates its validity as a construct. The latter is also reinforced by the fact that we were able to replicate this finding with a separate dataset.

Secondly, it is important to observe that different complexity measures, though correlated to each other and related to the same underlying construct of overall complexity, are nevertheless sensitive to different aspects of the data. In particular, measures operating along the temporal dimension appeared especially sensitive at discriminating between levels of sedation; conversely, topological measures failed to discriminate between Awake and Mild conditions in Dataset A, and also did not correlate with propofol serum levels. This suggests that the temporal dimension of the human brain’s complexity, as derived from BOLD signal timeseries (despite their limited temporal resolution compared to EEG), may be especially vulnerable to loss of consciousness, at least as it is induced by the GABA-ergic agent propofol. Further work may seek to identify whether this effect is uniform across cortical regions, or whether specific areas’ timeseries are more largely affected by propofol than others. This represents a novel insight regarding the ways in which anaesthetic drugs such as propofol intervene on the brain to cause unconsciousness. Additionally, it would be worth exploring whether this observation of different sensitivity of temporal and topological measures of complexity is drug-specific, or if instead it is a generalisable feature of how the brain loses consciousness. Thus, one future direction of research is to apply these same metrics to states of consciousness induced by different anaesthetic agents, whose molecular mechanisms of action can vary widely. Disorders of consciousness (DOC) due to severe brain injury may also represent a crucial future step for research: unlike anaesthetics, DOC involve changes in the physical structure of the brain, which is bound to impact the topology of brain networks. Investigating how this impacts the relation between different measures and dimensions of complexity will provide further understanding into the relation between complexity and consciousness in the brain. Additionally, as MRI is already a routine part of care for DOC patients, algorithms such as those explored here might be helpful in determining the presence or absence of consciousness in ambiguous states such as minimally conscious state.

Importantly, our results also show that, despite the relative temporal paucity of information in BOLD signals, these signals carry sufficient information to discriminate between states of consciousness. While preliminary, these findings suggest that the process complexity of individual BOLD signals is at least partially re-encoded as topological complexity when forming functional connectivity networks. One possible avenue of future work is to explore the parameters under which this conservation of complexity is maximized (different similarity functions, different thresholding procedures, etc), in order to increase the sensitivity of these measures. Crucially, even higher discriminative power may be achieved by applying the same analyses to measures with higher temporal information, such as EEG, which may then improve anaesthetists’ ability to detect unwanted residual consciousness in patients, thereby avoiding the rare but extremely distressing condition known as intraoperative awareness [27].

Nevertheless, our work also presents a number of limitations, and these should be borne in mind when evaluating the present results. Firstly, as already mentioned the temporal information available in the BOLD signal is limited, and it is also not a direct measure of neural activity. Additionally, our analysis pipeline involved removing the negative correlations between brain regions, before the network analysis. While negative correlations are unclear in origin and interpretation, and removing them is the most common approach, it is known that they are altered during anaesthesia and other states of unconsciousness [28, 29, 30]; thus, ignoring them may have different effects on conscious versus unconscious brain networks, which could explain the reduced sensitivity of topological measures. Thirdly, in Dataset A the state of consciousness was determined based on the estimated propofol concentration, rather than behaviour, so that different individuals’ susceptibility to the drug may have led to different levels of sedation, despite the same level of propofol. However, this concern is mitigated by the replication of our results in Dataset B, where sedation was deeper and it was assessed behaviourally, so that all individuals met the same criteria. Finally, the measures of complexity explored here are but a subset of those that have been proposed over the years in the literature. Future research could benefit from expanding this repertoire, for instance including estimates of Phi, a measure of integrated information derived from neural complexity [10], which has been proposed to quantify a system’s consciousness [9, 31]

## 5 Conclusion

We have investigated measures of algorithmic and process complexity of fMRI BOLD signal in both the temporal and topological dimensions, at various levels of consciousness induced by propofol sedation. Our results demonstrate that complexity measures are differently able to discriminate between levels of sedation, with temporal measures showing higher sensitivity. Additionally, all measures were strongly correlated, and most of the variance could be explained by a single underlying construct, which may be interpreted as a more general quantification of complexity, and which also proved capable of discriminating between levels of sedation, demonstrating a relation between consciousness and complexity.

## Supporting information

Final Data A

Final Data B

## Acknowledgements

This work was supported by a grant from the Wellcome Trust: Clinical Research Training Fellowship to Ram Adapa (Contract grant number: 083660/Z/07/Z); from the Canada Excellence Research Chairs program (215063) and the Canadian Institute for Advanced research (CIFAR)[to AMO]; Cambridge Biomedical Research Centre and NIHR Senior Investigator Awards [to DKM], the Stephen Erskine Fellowship at Queens College, Cambridge [to EAS], the LOral-Unesco for Women in Science Excellence Research Fellowship [to LN]; the British Oxygen Professorship of the Royal College of Anaesthetists [to DKM] and the Gates Cambridge Trust [to AIL]. I Pappas received funding from the Oon Khye Beng Ch’Hia Tsio Studentship for Research in Preventive Medicine, administered via Downing College, University of Cambridge; The research was also supported by the NIHR Brain Injury Healthcare Technology Co-operative based at Cambridge University Hospitals NHS Foundation Trust and University of Cambridge. We would like to thank Victoria Lupson and the staff in the Wolfson Brain Imaging Centre (WBIC) at Addenbrookes Hospital for their assistance in scanning. We would like to thank Dian Lu and Olaf Sporns for useful discussions, and all the participants for their contribution to this study.

## Footnotes

1 http://www.nitrc.org/projects/conn

2 https://github.com/spyder-ide/spyder

3 https://www.anaconda.com/download

